# Modified recipe to inhibit GSK-3 for the living fungal biomaterial manufacture

**DOI:** 10.1101/496265

**Authors:** Jinhui Chang, Po Lam Chan, Yichun Xie, Man Kit Cheung, Ka Lee Ma, Hoi Shan Kwan

**Author notes:** These authors contributed equally to this work (Jinhui Chang, Po Lam Chan).

## Abstract

Living fungal mycelium with suppressed or abolished fruit-forming ability is a self-healing substance particularly valuable biomaterial for further engineering and development in applications such as monitoring/sensing environmental changes and secreting signals. The ability to suppress fungal fruiting is also a useful tool for maintaining stability (*e.g.*, shape, form) of a mycelium-based biomaterial with ease and lower cost.

The objective of this present study is to provide a biochemical solution to regulate the fruiting body formation to replace heat killing of mycelium during production. We discovered that GSK-3 activity directly correlates with the development of fruiting bodies in fungi, especially mushroom forming fungi such as *Coprinopsis cinerea*. By regulating GSK-3 expression and activity, one can control the fungal fruiting body development.

We successfully demonstrated that treatment of an inhibitor of GSK-3 kinase activity resulted in acceleration in mycelium growth rate, absence of fruiting body and general decrease in GSK-3 gene expression. Therefore, GSK-3 inhibitor is suggested to be included in the mycelium cultivation recipes for regulating the growth of fungal mycelium and for inhibiting the development of fruiting bodies. This is the first report of using a GSK-3 inhibitor, such as lithium or any other GSK-3 inhibitor, to suppress or abolish fruiting body formation in living fungal mycelium-based biomaterial. It also provides an innovative strategy for easy, reliable, and low cost maintenance of biomaterial containing live fungal mycelium.

## Introduction

Fungal mycelium-based biomaterials are fast emerging in recent years. Mycelium is the vegetative structure of fungi, mainly composed of natural polymers. Mycelium-based biomaterials have a wide range of applications due to their controlled and tunable properties during growth, its self-assembly, self-healing, environmentally responsive, and biodegradable nature. The dried mycelium has strength, durability, and many other beneficial qualities: it is nontoxic, fire-resistant, mold resistant, water-resistant, and a great thermal insulator, amongst other salient features(1–13). Under proper circumstances, mycelium of many mushroom-forming fungi will aggregate to form mushrooms, which are the fruiting body spreading spores(14). Not only the fruiting bodies will cause conformational changes of the mycelium based materials, but also the spores may cause allergy and infection in susceptible population. In current production procedures of the mycelium-based materials, whole material are heated or treated with fungicide to kill the living cells, to stop the fruiting body formation(15). Such rendered mycelium-based materials retain few of the benefits of living material. Therefore, new approaches are needed for inhibiting fruiting body formation while keeping the mycelium alive, in order to produce living mycelium-based materials of desirable qualities.

Kinases mediate cellular and developmental responses to growth factors, environmental signals, and internal processes, and the kinases cascades play crucial roles in many signaling transduction pathways(16,17). Phosphorylation of proteins kinases affects their activity, localization, stability, conformation, and protein-protein interaction. One interesting and putatively central regulatory kinase is glycogen synthase kinase-3 (GSK-3). GSK-3 is a serine/ threonine kinase of the CMGC family of proline-directed kinases that is highly conserved in all eukaryotes. GSK-3 is activated by the constitutive phosphorylation at a C-terminal tyrosine residue, however, the regulatory phosphorylation at an N-terminal serine residue causes a conformational change to block the catalytic domain, hence inhibits its kinase activity (18). The kinases PKA, PKB, and PKC inhibit GSK-3 in specific signaling pathways in eukaryotes, while in fungi these kinases are essential growth regulators in response to environmental stimuli(19–21).

In mammals, GSK-3 inhibition has attracted widespread attention as one of the critical therapeutic targets whereby lithium exerts its pharmacological effects on mood stabilization, neurogenesis, neurotrophicity, neuroprotection, anti-inflammation, and others(18). Lithium compounds are also suggested to be added in cultivation to fortify the lithium nutrient value of some edible mushrooms (22,23). Lithium chloride (LiCl) is a well-known substance that has been shown to inhibit GSK-3 and recent evidence suggests that low, non-toxic concentrations of such a compound have indeed anti-inflammatory effects(24).

In this study, *Coprinopsis cinerea* is used to represent the white-rot basidiomycetous fungi, as it is a classic model mushroom-forming fungus(25). The typical life cycle of *C. cinerea* can be finished within 2 weeks under lab condition, which includes stages of basidiospores, vegetative mycelium, hyphal knots, initial, stage-1 and -2 primordia, young and mature fruiting bodies(26).

This study aims to provide a biochemical approach to inhibit fruiting body formation from the mycelium-based biomaterals, thus producing living biomaterials. We demonstrated that fruiting body development in mushrooms can be regulated by modulating GSK-3 expression and/or activity: suppression of GSK-3 expression and/or activity can promote the growth of mycelium and inhibit the fruiting body formation, whereas enhancement of GSK-3 expression and/or activity can achieve opposite effects. Regulation of GSK-3 can be applied in the manufacturing of mycelium-based biomaterials, which can shorten the production cycle, reduce the cost for maintenance of mycelium materials, and therefore achieve a higher level of cost-effectiveness.

## Materials and Methods

### Strains and cultivation conditions

Two GSK3 inhibitors (LiCl and CHIR99021-HCl) and one GSK3 activator (Cisplatin) were tested in *Coprinopsis cinerea*, the homokaryotic fruiting strain #326 (*A43mut B43mutpab1-1*). One GSK3 inhibitor (LiCl) was tested in *Pleurotus djamor* (commonly known as the pink oyster mushroom). Belonging to the same order Agaricales, these two tested mushroom species are of two different families, Psathyrellaceae and Pleurotaceae, respectively. *C. cinerea* is cultured on yeast extract-malt extract-glucose (YMG) agar plates (per litre, 4 g yeast extract; 10 g malt extract; 4 g glucose; 10 g agar;) at 37°C in the dark until mycelia grow over the whole agar surface (27). Fruiting body development is induced by incubating the mycelia at 25°C under a 12hours light /12hours dark cycle. *P. djamor is* cultivated on Potato Dextrose Agar (PDA, BD Difco™) plates at 28 °C in the dark until mycelia grow over the whole agar surface, and transferred to 25 °C 12hours light /12hours dark cycle to induce fruiting body formation. Triplicates were employed in each set up. Each plate was measured for 45g (±1g) medium to uniform the nutrients contents and for accurate inhibitor concentration.

### Effect of Inhibitors and Activator

Inhibitors of GSK-3 used in this study include lithium chloride (LiCl), as well as CHIR99021 trihydrochloride. Three methods have been tested to deliver LiCl (Sigma-Aldrich, St. Louis, MO, USA). One delivery method is to mix 1.5g/L, 3 g/L or 6g/L LiCl in the medium before autoclave sterilization, and the other methods are to spread LiCl solution on the surface of agar before inoculation, or add LiCl solution under the agar after mycelia reaches the petri dish edge. CHIR99021 trihydrochloride is very specific and water soluble, so it is used to confirm the effect of inhibited GSK-3 on fruiting body development. 5nM, 10nM and 15nM of CHIR99021 trihydrochloride is spread on the surface of agar evenly before inoculation.

In this study, one specific GSK-3 activator, cisplatin, is tested. It is also known as cisplatinum, platamin, neoplatin, cismaplat, or cis-diamminedichloridoplatinum(II) (CDDP), a chemical most commonly used in chemotherapy in cancer treatment. 1 ml Water or saturated Cisplatin solution (25 °C) was filtered and spread on the surface of YMG agar, before inoculation. CHIR99021 trihydrochloride and cisplatin are not suggested to be autoclaved, so we use 0.2 micron filter to remove all bacteria in the solution.

### Sensitive window to LiCl

The effect of LiCl at different developmental stages of *C. cinerea* are tested to find the sensitive window. 0.1ml of 3g/L LiCl or 0.1 ml water were added under the agar at the stages of: mycelium, hyphal knot, initial, stage-1primordium, stage-2 primordium and young fruiting body. The growth status was record till 3 days after the control group form mature fruiting bodies.

### Expression levels of GSK-3 downstream target genes

The expression levels of targets genes of GSK-3 were measured to explore the mechanism of LiCl. The GSK-3 downstream targets were predicted by Orthologue comparison. A total of 83 GSK-3 downstream targets reported in human and mouse, were compared to the *C. cinerea* genes by OrthoMCL V2.0 (28), and 52 orthologues were identified. Among them, glycogen synthase (CC1G_01973), GSK-3 (CC1G_03802), eukaryotic translation initiation factor 1 (eIF1) (CC1G_03881), uncharacterized protein with Ricin B-type lectin domain (CC1G_05298), translation initiation factor eIF2 gamma subunit (CC1G_09429) were picked for real-time PCR analysis. For each gene, primers for two segments with similar PCR condition were selected.

Samples treated by water or 1.5g/L or 3g/L LiCl (mixed before autoclave) were cultured as conditions described above. Once the control group started fruiting body initiation, total RNAs of three biological replicates were extracted using RNeasy^®^ Plant Mini Kit (Qiagen). RNA products were stored at −80°C. The concentration of RNA was measured by NanoDrop Spectrophotometers (Thermo Scientific). 500ng RNA products were used to synthesize cDNA using iScript TM cDNA Synthesis Kit (Bio-Rad). Quantitative real-time PCR (qPCR) was performed by Applied Biosystems^®^ 7500/7500 Fast Real-Time PCR system™ (Applied Biosystems) using iQTM SYBR^®^ Green Supermix (Bio-Rad).

## Results

### Effect of GSK-3 inhibitors and activator on fruiting body development

As shown in Figure 1a, effect of LiCl to *C. cinerea* fruiting body development was tested. While the control group has already developed into mature fruiting bodies, primordium, initials and hyphal knots were formed on the plates treated with 1.5g/L LiCl respectively. The plates treated with 3g/L LiCl and 6g/L LiCl were arrested in mycelium stage, and mycelium treated with 6g/L LiCl grew than other groups. These results showed that LiCl of higher concentrations have stronger inhibitory effect on *C. cinerea* fruiting body development.

The three delivery methods of LiCl, either mixed in the agar, on the surface of agar, or under the agar, show no differences and all can efficiently inhibit the fruiting body development. LiCl is not sensitive to heat treatment, and can be autoclaved to sterilize the solution. Any of these delivery methods can be chosen in a large-scale manufacture of the biomaterials.

In order to confirm that the inhibition of fruiting body development is caused by GSK-3 inhibition, another more specific GSK-3 inhibitor, CHIR99021 trihydrochloride, was tested. As shown in 1b,Young fruiting bodies developed on the control plates treated with water and the plates with 1 μM CHIR99021 trihydrochloride. The plates treated with 100 μM CHIR99021 trihydrochloride developed primordium. The plates treated with 500 μM CHIR99021 trihydrochloride remained arrested in mycelium stage. These results showed an stronger inhibitory effect on *C. cinerea* fruiting body development by CHIR99021 trihydrochloride at higher concentrations.

LiCl and CHIR99021 trihydrochloride treatments inhibited the fruiting body initiation in a dose-dependent manner (Figure 1a and 1b). Given that both LiCl and CHIR99021 trihydrochloride are specific inhibitors to GSK-3, it can be concluded that the effect of lithium is mediated through the inhibition of GSK-3 activity.

To demonstrate that GSK-3 is also important in fruiting body formation in other mushrooms, the effect of LiCl to *Pleurotus djamor* fruiting body development was tested (Figure 1c). *C. cinerea* and *P. djamor* are two fungal species within the same family of Basidiomycota and the same order of Agaricales.

LiCl was added to YMG agar medium before autoclave. Mature fruiting bodies developed on the control plates. The plates treated with 2g/L LiCl failed to develop fruiting body in the following 30 days. These results showed an inhibitory effect on *P. djamor* fruiting body development by LiCl.

With the positive results that GSK-3 inhibitor can inhibit the fruiting body development, it is hypothesized that GSK-3 activity is associated with the fruiting body development. Then a GSK-3 activator, cisplatin, was tested for its effect on *C. cinerea* fruiting body development (Figure 1d). The activator treated group with 1 ml saturated Cisplatin, showed an accelerated development since the formation of hyphal knot, and the mature fruiting body appeared 2 days earlier than the control group, while the control group was slower and only developed into young fruiting body. These results showed a positive or promoting effect on *C. cinerea* fruiting body development by GSK-3 activator, Cisplatin.

**Fig. 1.**
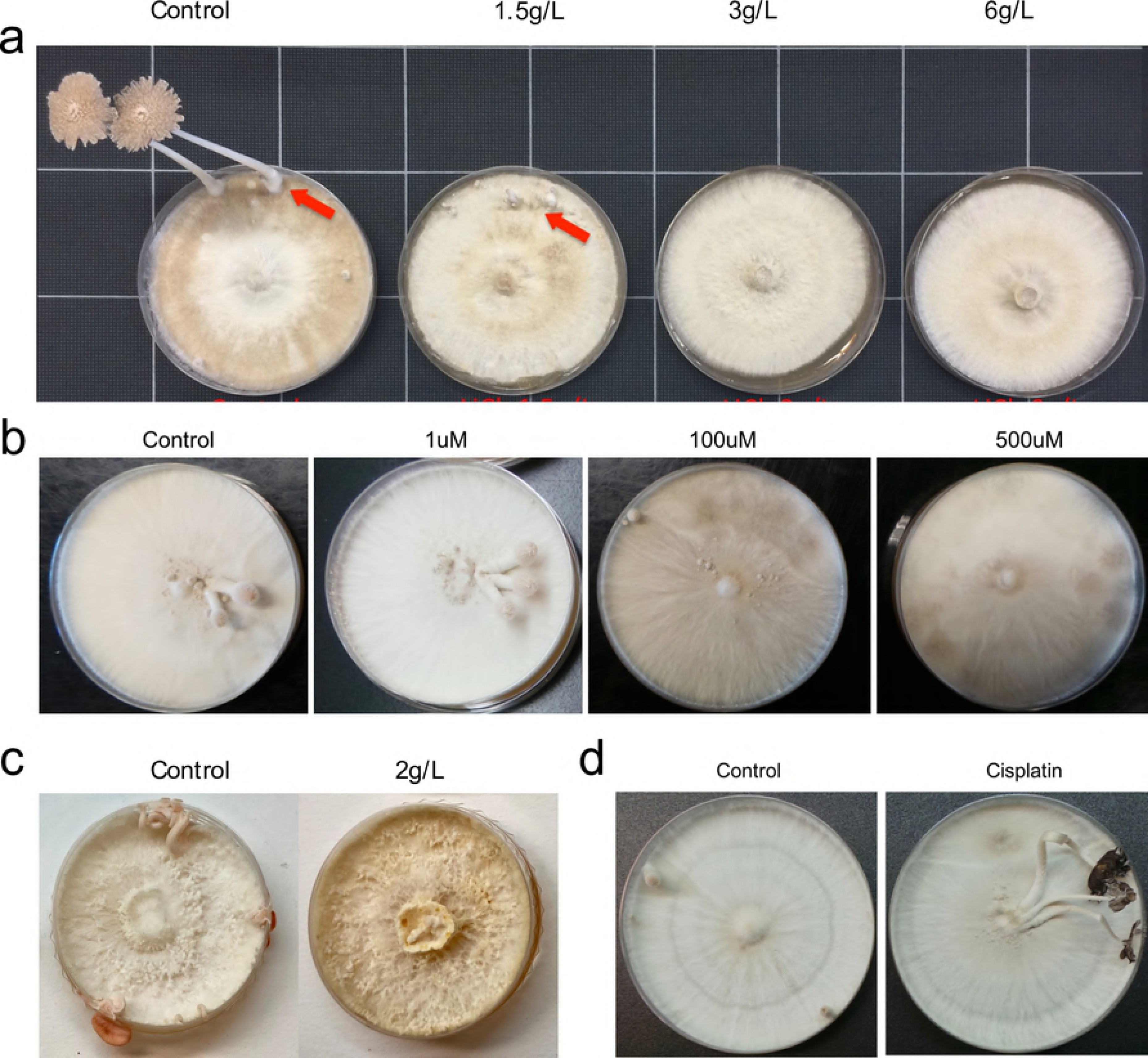
Effect of GSK-3 inhibitors and activator on the fruiting body development. **(a)**The different doses of GSK-3 inhibitor, LiCl, have different levels of inhibitory effect on the development of *C. cinerea*. Mature fruiting bodies, young fruiting bodies, primordia and initials were producd in the control group, while the 1.5g/L LiCl treated group produced only initials and primordia. No initiation was observed in the groups treated with higher concentration of LiCl, in the following 30 days. **(b)** The different doses of GSK-3 inhibitor, CHIR99021 trihydrochloride, have different levels of inhibitory effect on the development of *C. cinerea*. Young fruiting bodies developed in the control group and 1 μM treated group, while only primordium were developed on plates treated with 100 μM. The plates treated with 500 μM remained in mycelium stage in the following 30 days. **(c)** GSK-3 inhibitor, LiCl, also inhibit he development of *P. djamor*. Mature fruiting bodies, young fruiting bodies, primordia and initials were produced in the control group, while the 2g/L LiCl remained in mycelium stage in the following 30 days. **(d)** GSK-3 activator, cisplatin, accelerates the development of *C. cinerea*. After 6-days incubation, the plates treated with 1 ml saturated Cisplatin solution had fruiting body and began autolysis, while the control group was slower and only developed into young fruiting body.

These data unequivocally support the conclusion that among fungus species of the division Basidiomycota, especially of the order Agaricales, GSK3 inhibitors inhibit/reduce/decelerate the fruiting body formation, whereas GSK3 enhancers activate/increase/accelerate the fruiting body formation.

### GSK-3 inhibitor promotes mycelium growth

The mycelial growth rate was measured for the LiCl treated *C. cinerea*. Biological triplicates were cultivated on YMD at 38°C in darkness, treated with different doses of LiCl. The mycelium area was recorded for 4 days. Figure 2a shows the average growth area of each group. As the inoculum usually need time to adapt to the new environment and absorb nutrients, there was no differences on the first two days. The differences apear on the 3^rd^ day, the LiCl treated groups spanned faster than the control group, and after 144 hours of inoculation the mycelium spanned into a circle shape of 51.8 cm^2^ for 1.5g/L, and 55.1 cm^2^ for 3 g/L, while 42.9 cm^2^ for control group. The results show that proper concentrations of LiCl can accelerate the mycelium growth.

This is an ideal property of GSK-3 inhibitor for the large-scale manufacture of mycelium-based biomaterial. The modified recipe with addition of porper concentration of LiCl, can not only inhibit the appearance of fruiting body, but also speed up the mycelium growth, and hence lower the shorten the manufacture cycle and lower the cost.

### Effect of GSK-3 inhibitor on gene expression levels

To validate the LiCl is targeting on GSK-3 other than other gene, we test the expression levels of the GSK-3 gene and its downstream target genes. Two segments of each gene were tested for higher accuracy. The real-time PCR results show that the LiCl could regulate the expression of the GSK-3 gene itself, as well as the selected GSK-3 target genes. GSK-3, glycogen synthase, and protein with Ricin B-type lectin domain showed decrease in gene expression levels with increase of LiCl concentration. eukaryotic translation initiation factor 1 is stable in all conditions, while translation initiation factor eIF2 gamma subunit increase to 2-fold in 1.5g/L LiCl treated group. The change in gene expression supports that LiCl targets the GSK-3 and affects its downstream genes.

**Fig. 2.**
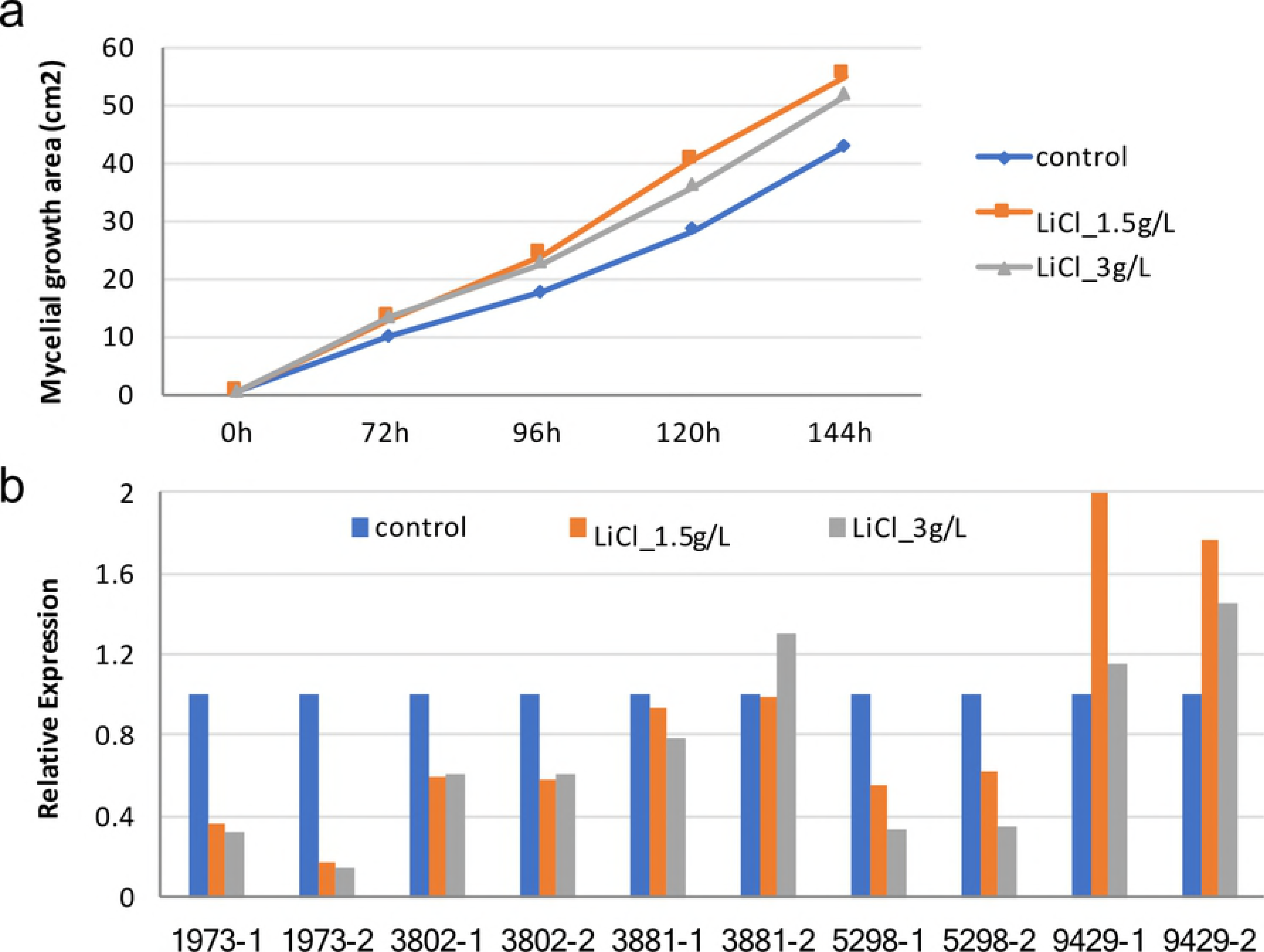
**(a)** Mycelial growth of *C. cinerea* with different doses of LiCl, biological triplicates were used for mean calculation. The growth rate of mycelium treated with 1.5g/L and 3g/L LiCl is higher than control. **(b)**The gene expression levels are indicated by the abundance of segments on the cDNA, by real-time PCR. The GSK-3 and its target genes change the expression level under LiCl treatment. 1973-1 & 1973-2 are two segments on the cDNA of glycogen synthase (CC1G_01973), and the others are segments of the following transcripts: GSK-3 (CC1G_03802); eukaryotic translation initiation factor 1 (CC1G_03881); Uncharacterized protein with Ricin B-type lectin domain (CC1G_05298); Translation initiation factor eIF2 gamma subunit (CC1G_09429).

### Sensitive Window to GSK-3 inhibitor LiCl

For the current manufacture of mycelium-base biomaterial, it’s difficult to avoid the formation of fruiting body. Upon environmental stimuli, including nutrient depletion, light/dark cycle, and cold shock, mycelia aggregate into hyphal knot, followed by fruiting body initials. Initials then develop into stage-1 and -2 primordia, young and mature fruiting bodies. The companies may already have fixed production line, so we want to explore all the possible procedures to introduce the GSK-3 inhibitor, specifically, LiCl, which is cheaper compared to other GSK-3 inhibitors.

In the previous sections, LiCl is demonstrated to arrest the mycelium. Hyphal knot is a short stage which is difficult to define by naked eyes. So we only test the stages after initials. As shown in Figure 3a, adding 3g/L LiCl at stages of initial and stage-1primordium led to the arrest of their development, while stage-2 primordium and young fruiting body could continue to develop into mature fruiting bodies. Intervention of LiCl at stages of mycelium, initial and stage-1 primordium resulted in arrestment in fruiting body development, so these stages are sensitive windows to LiCl.

**Fig. 3.**
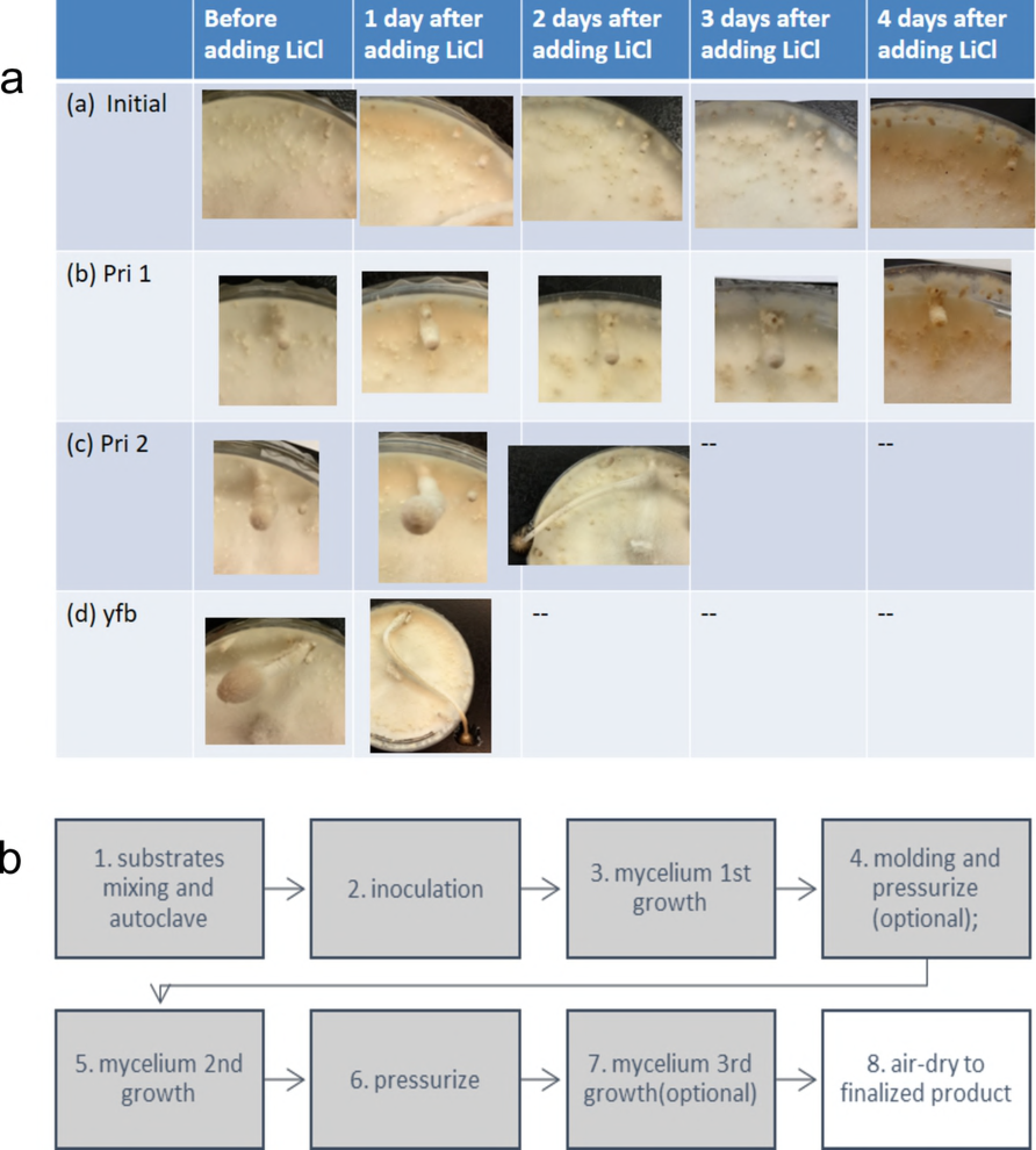
**(a)**The development of *C. cinerea* treated with 0.1ml of 3g/L LiCl at different stage. Adding 3g/L LiCl at stages of initial and stage-1 primordium led to inhibition on further development. Adding 3g/L LiCl at stages of stage-2 primordium and young fruiting body cannot inhibit the fruiting body development. **(b)** Proposed procedures of the production pipeline of living mycelium-based materials, and the GSK-3 inhibitors can be added at any time from procedure 1 to 7.

For producing living mycelium-based material, the basic production pipeline is designed for adding GSK-3 inhibitors, particularly the lithium or lithium salt. The production pipeline can be all of part of the following procedures: 1) substrates mixing and autoclave; 2) inoculation; 3) mycelium 1st growth; 4) molding and pressurize; 5) mycelium 2nd growth; 6) pressurize (optional); 7) mycelium 3rd growth (optional); 8) air-dry to finalized product. LiCl or other GSK-3 inhibitors can be added at any time from procedure 1) to 7), by mixing in the substrate before autoclave, adding the sterilized solution on the surface or inside the medium mixture before inoculation, or spraying to the mycelium after a period of growth.

### Discussion

The study demonstrated that GSK-3 activity can determine the fruiting body development, and GSK-3 inhibitor is suggested to be included into recipes to manufacture live fungal mycelium that exhibits an altered and more desirable profile of fruiting body development as well as compositions that contain the fungal mycelium.

In many instances one would prefer for a live fungal mycelium to refrain from developing fruiting bodies such that the mycelium is easily maintained without concerns of loss in its shape, form, or consistency. In contrast to the currently-in-use method of heat-killing fungal mycelium to prevent fruiting body formation, a live version of mycelium that simply does not form fruiting bodies is far more desirable considering its live nature and thus healing potential.

In other cases, promoting fruiting body development may be of interest. For instance, when the intended goal is to produce and harvest as many fruiting bodies (*e.g.*, mushrooms and truffles) as possible in a defined time period, having enhanced fruiting body development is beneficial.

Glycogen synthase kinase-3, GSK-3 has important role in cell-fate specification, leading to cell differentiation or apoptosis or development through number of signaling pathways(29–32). So we propose a pathway that GSK-3 could be the links between environmental stimuli and the responsive development, and a master-switch of fruiting body formation (Figure 4).

**Fig. 4.**
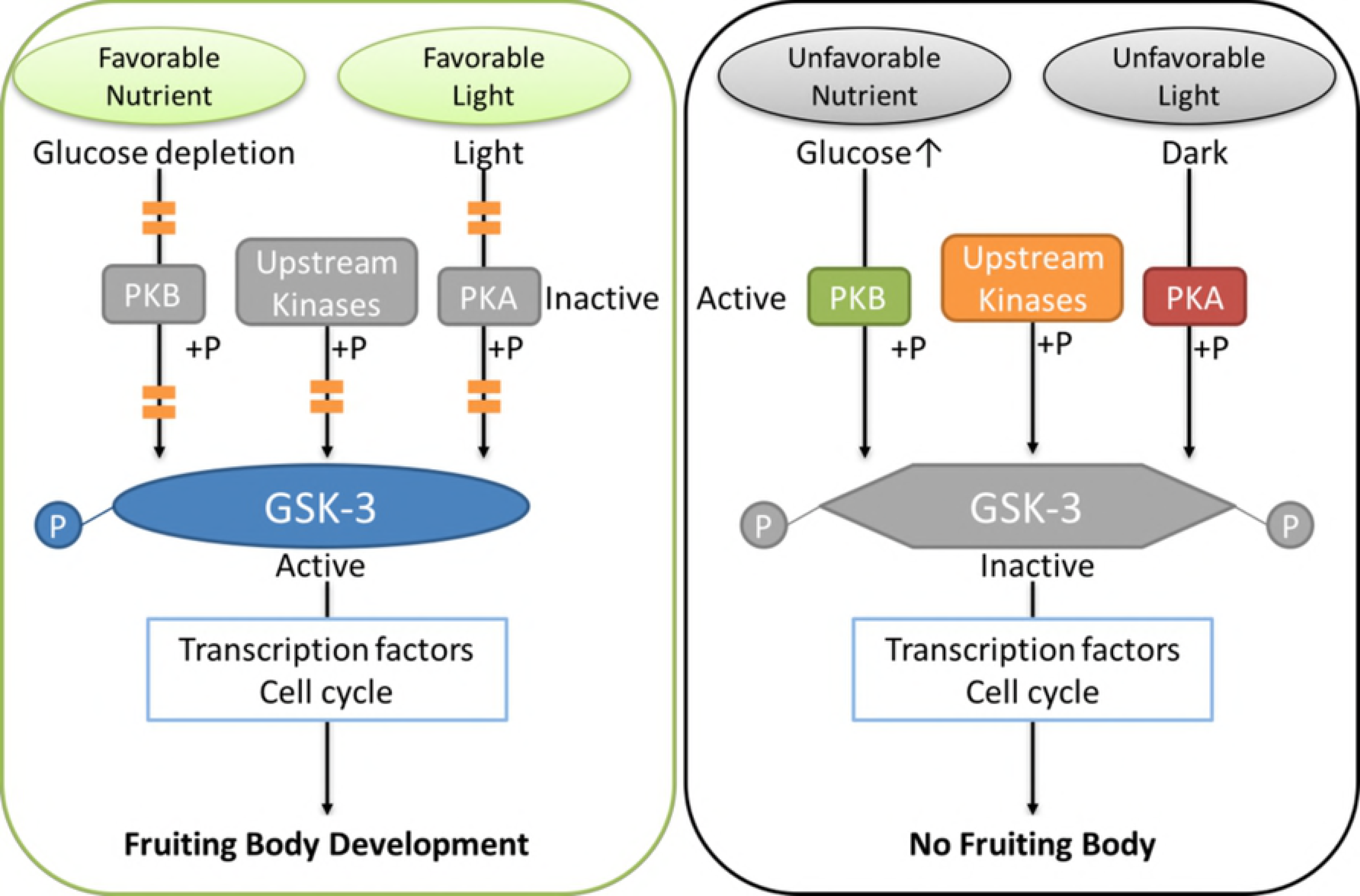
Glycogen synthase kinase-3 (GSK-3) as a master-switch of fruiting body formation. GSK-3 has important role in cell-fate specification, leading to cell differentiation or apoptosis or development through number of signaling pathways. GSK-3 could be the links between environmental stimuli and the responsive development, and a master-switch of fruiting body formation. The activity of GSK-3 determines the fruiting body development.

For producing a live fungal mycelium with an enhanced or inhibited fruiting body development profile, either a permanent means (*e.g.*, GSK-3 knockdown or GSK-3 knockout fungal strain) or a transient means (*e.g.*, application of an activator or inhibitor of GSK-3 present in the medium for fungi) can be employed. While the former may be easier to maintain in the long term, efforts involved in the initial stage of establishing the genetically modified fungal strains are tremendously more significant both in cost and in time. In contrast, the latter offers the benefits of flexibility and low-cost use, when the GSK-3 activator or inhibitor can be readily removed at an appropriate time such that the fungus may resume its normal life cycle of different phases.

One possibility is to reduce or abolish GSK-3 expression by genetic manipulation of the fungal cells’ genomic sequence encoding the GSK-3 protein or by transient or permanent expression of small inhibitory RNAs. The gene encoding GSK-3 is highly conserved across diverse phyla. It is a serine/threonine kinase having a monomeric structure and a size of approximately 47 kilo daltons. The amino acid sequence and corresponding polynucleotide coding sequence for *C. cinerea* GSK-3 are provided in GenBank Accession Numbers XP_001833585 (strain okayama7#130) and NW_003307543.1 (Genomic Sequence in strain Okayama7#130) / jgi|Copci_AmutBmut1|363162|e_gw1.29.187.1 (in strain #326, Taxonomy ID: 1132390 and Accession: PRJNA258994), respectively. The GSK-3 is highly conserved in protein sequence. The homologous proteins include PIL30457.1 in *Ganoderma sinense* ZZ0214-1, and jgi|Gansp1|158466|gm1.11165_g in Ganoderma sp. 10597 SS1 (North American isolate of *G. lucidum*), and KDQ33621 in *Pleurotus ostreatus* PC15. As used herein, a GSK-3 protein encompasses both *C. cinerea* GSK-3 protein and its homologs/orthologs in fungal species, especially those of Basidiomycota, having at least 90% or more sequence homology to *C. cinerea* GSK-3 protein sequence and share essentially the same biological or enzymatic activity.

A GSK-3 knockdown cell may be generated by genetic manipulation of the genomic GSK-3 sequence of a suitable parent cell. Methods such as sequence homology-based gene disruption methods utilizing a viral vector or CRISPR system can be used for altering the GSK-3 genomic sequence, for example, by insertion, deletion, or substitution, which may occur in the coding region of the gene or in the non-coding regions (*e.g.*, promoter region or other regulatory regions) and which may result in complete abolition of GSK-3 expression, reduced GSK-3 expression, or unaltered expression at mRNA level but diminished GSK-3 protein activity.

Another possibility is to suppress the activity of endogenously expressed GSK-3 protein by introducing a GSK-3 inhibitor into the external environment in which the fungi grow. Lithium salts have mycelium-enhancing effect to some mushroom forming fungi, but the concentration range of such effect is narrow. In some other mushroom forming fungi, high concentration of LiCl may inhibit the mycelium growth, especially in *Trichoderma* species, which is a common contamination of the edible mushroom (33). Thus, while LiCl might be applied to prevent fruiting in some mushroom-forming fungi, it can also inhibit the contamination during manufacturing in some scenario. In addition to LiCl, other agents that specifically target GSK-3, can also prevent the development of fruiting body. In support of this conclusion, CHIR99021 trihydrochloride, an alternative GSK-3 specific inhibitor that acts through a distinct mechanism, also inhibits fruiting body formation. Other known GSK-3 inhibitors include: Maleimide Derivatives; Staurosporine and Organometallic Inhibitors; Indole Derivatives; Paullone Derivatives; Pyrazolamide Derivatives; Pyrimidine and Furopyrimidine Derivatives; Oxadiazole Derivatives; Thiazole Derivatives; and Miscellaneous Heterocyclic Derivatives. (18,34)

The sensitive window to LiCl is from mycelium, hyphal knot, initial to stage-1 primordium. This indicates that the LiCl may inhibit fruiting through affecting the cell differentiation. Inhibitors of GSK-3 were shown to maintain the mouse and human embryonic stem (ES) cells in undifferentiated status, while removing inhibitor promotes differentiation into multiple cell lineages (35). The potency maintaining function of GSK-3 may be related to protein degradation. After phosphorylated by GSK-3, many substrates will then be targeted by ubiquitination for proteasome-mediated degradation. Undifferentiated cells are proliferative because GSK-3 activity is limited by persistent unfavorable growing condition signals. The effectors of GSK-3, such as transcription factors, are less modified by phosphorylation and ubiquitination, so their half-lives are prolonged to enhance stem/precursor cell proliferation (36). This suggested GSK3 interferes fruiting by interfering cell specification. Deeper studies are needed to discover the mechanism in detail.

## Reference

1. Jones M, Bhat T, Huynh T, Kandare E, Yuen R, Wang CH, et al. Waste-derived low-cost mycelium composite construction materials with improved fire safety. Fire and Materials. 2018 Nov;42(7):816–25.

2. Nguyen PQ, Courchesne NMD, Duraj-Thatte A, Praveschotinunt P, Joshi NS. Engineered Living Materials: Prospects and Challenges for Using Biological Systems to Direct the Assembly of Smart Materials. Vol. 30, Advanced Materials. 2018. p. 1704847.

3. Karana E, Blauwhoff D, Hultink E, Camere S. When the Material Grows : A Case Study on Designing (with) Mycelium-based Materials. 2018;12(2):119–36.

4. Nguyen PQ, Courchesne NMD, Duraj-Thatte A, Praveschotinunt P, Joshi NS. Engineered Living Materials: Prospects and Challenges for Using Biological Systems to Direct the Assembly of Smart Materials. Vol. 30, Advanced Materials. 2018.

5. Grimm D, Wösten HAB. Mushroom cultivation in the circular economy. Vol. 102, Applied Microbiology and Biotechnology. 2018. p. 7795–803.

6. Appels FVW, Dijksterhuis J, Lukasiewicz CE, Jansen KMB, Wösten HAB, Krijgsheld P. Hydrophobin gene deletion and environmental growth conditions impact mechanical properties of mycelium by affecting the density of the material. Scientific Reports. 2018;8(1).

7. Camere S, Karana E. Fabricating materials from living organisms: An emerging design practice. JOURNAL OF CLEANER PRODUCTION. 2018;186:570–84.

8. Khamrai M, Banerjee SL, Kundu PP. A sustainable production method of mycelium biomass using an isolated fungal strain Phanerochaete chrysosporium (accession no: KY593186): Its exploitation in wound healing patch formation. Biocatalysis and Agricultural Biotechnology. 2018;16:548–57.

9. Jones M, Bhat T, Huynh T, Kandare E, Yuen R, Wang CH, et al. Waste-derived low-cost mycelium composite construction materials with improved fire safety. Fire and Materials. 2018;

10. Kilaru S, Hoegger PJ, Kües U. The laccase multi-gene family in Coprinopsis cinerea has seventeen different members that divide into two distinct subfamilies. Current Genetics. 2006 Jul 28;50(1):45–60.

11. Grimm D, Wösten HAB. Mushroom cultivation in the circular economy. Vol. 102, Applied Microbiology and Biotechnology. 2018. p. 7795–803.

12. Appels FVW, Camere S, Montalti M, Karana E, Jansen KMB, Dijksterhuis J, et al. Fabrication factors influencing mechanical, moisture- and water-related properties of mycelium-based composites. Materials & Design. 2018;

13. Silverman J. Development and testing of mycelium-based composite materials for shoe sole applications. 2018;

14. Kües U, Subba S, Yu Y, Sen M. Regulation of fruiting body development in Coprinopsis cinerea. 2016;(July).

15. Haneef M, Ceseracciu L, Canale C, Bayer IS, Heredia-Guerrero JA, Athanassiou A. Advanced Materials From Fungal Mycelium: Fabrication and Tuning of Physical Properties. Scientific Reports. Nature Publishing Group; 2017;7(January):41292.

16. Kosti I, Mandel-Gutfreund Y, Glaser F, Horwitz B. Comparative analysis of fungal protein kinases and associated domains. BMC Genomics. 2010 Feb;11(1):133.

17. Wang C, Zhang S, Hou R, Zhao Z, Zheng Q, Xu Q, et al. Functional analysis of the kinome of the wheat scab fungus Fusarium graminearum. Howlett BJ, editor. PLoS pathogens. Public Library of Science; 2011 Dec;7(12):e1002460.

18. Takahashi-yanaga F. Activator or inhibitor ? GSK-3 as a new drug target. Biochemical Pharmacology. Elsevier Inc.; 2013;86(2):191–9.

19. Andoh T, Hirata Y, Kikuchi A. Yeast glycogen synthase kinase 3 is involved in protein degradation in cooperation with Bul1, Bul2, and Rsp5. Molecular and cellular biology. 2000 Sep;20(18):6712–20.

20. Moore SF, van den Bosch MTJ, Hunter RW, Sakamoto K, Poole AW, Hers I. Dual regulation of glycogen synthase kinase 3 (GSK3)α/β by protein kinase C (PKC)α and Akt promotes thrombin-mediated integrin αIIbβ3 activation and granule secretion in platelets. The Journal of biological chemistry. 2013 Feb 8;288(6):3918–28.

21. Tataroğlu Ö, Lauinger L, Sancar G, Jakob K, Brunner M, Diernfellner ACR. Glycogen synthase kinase is a regulator of the circadian clock of Neurospora crassa. The Journal of biological chemistry. 2012 Oct 26;287(44):36936–43.

22. Mleczek M, Siwulski M, Rzymski P, Budzyńska S, Gąsecka M, Kalač P, et al. Cultivation of mushrooms for production of food biofortified with lithium. European Food Research and Technology. 2017;243(6):1097–104.

23. De Assunão LS, Da Luz JMR, Da Silva MDCS, Vieira PAF, Bazzolli DMS, Vanetti MCD, et al. Enrichment of mushrooms: An interesting strategy for the acquisition of lithium. Food Chemistry. Elsevier; 2012 Sep 15; 134(2): 1123–7.

24. Zhang F, Phiel CJ, Spece L, Gurvich N, Klein PS. Inhibitory phosphorylation of glycogen synthase kinase-3 (GSK-3) in response to lithium. Evidence for autoregulation of GSK-3. The Journal of biological chemistry. American Society for Biochemistry and Molecular Biology; 2003 Aug 29;278(35):33067–77.

25. Stajich JE, Wilke SK, Ahrén D, Hang C, Birren BW, Borodovsky M, et al. Insights into evolution of multicellular fungi from the assembled chromosomes of the mushroom Coprinopsis cinerea (Coprinus cinereus).

26. Plaza D, Lin C-W, van der Velden NS, Aebi M, Künzler M, Dacks J, et al. Comparative transcriptomics of the model mushroom Coprinopsis cinerea reveals tissue-specific armories and a conserved circuitry for sexual development. BMC Genomics. BioMed Central; 2014;15(1):492.

27. Rao PS, Niederpruem DJ. Carbohydrate metabolism during morphogenesis of Coprinus lagopus (sensu Buller). Journal of bacteriology. 1969 Dec;100(3):1222–8.

28. Li L, Stoeckert CJ, Roos DS. OrthoMCL: identification of ortholog groups for eukaryotic genomes. Genome research. Cold Spring Harbor Laboratory Press; 2003 Sep 1;13(9):2178–89.

29. Tataroglu O. Role of Glycogen Synthase Kinase (GSK) in temperature compensation of the. 2011;1–88.

30. Ninkovic J, Stigloher C, Lillesaar C, Bally-Cuif L. Gsk3beta/PKA and Gli1 regulate the maintenance of neural progenitors at the midbrain-hindbrain boundary in concert with E(Spl) factor activity. Development (Cambridge, England). 2008;135(18):3137–48.

31. Casas-Flores S, Rios-Momberg M, Rosales-Saavedra T, Martínez-Hernández P, Olmedo-Monfil V, Herrera-Estrella A. Cross Talk between a Fungal Blue-Light Perception System and the Cyclic AMP Signaling Pathway. Eukaryotic Cell. 2006 Mar;5(3):499–506.

32. Zanolli F, Magalhães R, Paula D, Carlos L, Barbosa B, Francisco H, et al. cAMP signaling pathway controls glycogen metabolism in Neurospora crassa by regulating the glycogen synthase gene expression and phosphorylation. Fungal Genetics and Biology. Elsevier Inc.; 2010;47(1):43–52.

33. H.G. Wildman. Lithium chloride as a selective inhibitor of Trichoderma species on soil isolation plates. Mycological Research. Elsevier; 1991 Dec 1;95(12):1364–8.

34. Selenica M-L, Jensen HS, Larsen AK, Pedersen ML, Helboe L, Leist M, et al. Efficacy of small-molecule glycogen synthase kinase-3 inhibitors in the postnatal rat model of tau hyperphosphorylation. British journal of pharmacology. Blackwell Publishing; 2007 Nov 12;152(6):959–79.

35. Kirby LA, Schott JT, Noble BL, Mendez DC, Caseley PS, Peterson SC, et al. Glycogen synthase kinase 3 (GSK3) inhibitor, SB-216763, promotes pluripotency in mouse embryonic stem cells. Cooney AJ, editor. PloS One. Public Library of Science; 2012 Jan;7(6):e39329.

36. Westermarck J. Regulation of transcription factor function by targeted protein degradation: an overview focusing on p53, c-Myc, and c-Jun. Methods in molecular biology (Clifton, NJ). 2010 Jan;647:31–6.

